# Single-cell RNA sequencing reveals a dynamic stromal niche within the evolving tumour microenvironment

**DOI:** 10.1101/467225

**Authors:** Sarah Davidson, Mirjana Efremova, Angela Riedel, Bidesh Mahata, Jhuma Pramanik, Jani Huuhtanen, Gozde Kar, Roser Vento-Tormo, Tzachi Hagai, Xi Chen, Muzlifah A. Haniffa, Jacqueline D. Shields, Sarah A. Teichmann

**Affiliations:** Wellcome Sanger Institute, Wellcome Genome Campus, Hinxton, Cambridge CB10 1SA, UK.; Medical Research Council Cancer Cell Unit, Hutchison/Medical Research Council Research Centre, Cambridge, UK.; Hematology Research Unit Helsinki, University of Helsinki, Finland.; Institute of Cellular Medicine, Newcastle University, Newcastle upon Tyne, NE2 4HH, UK

## Abstract

Non-cancerous stromal cells represent a highly diverse compartment of the tumour, yet their role across tumour evolution remains unclear. We employed single-cell RNA sequencing to determine stromal adaptations in murine melanoma at different points of tumour development. Naive lymphocytes recruited from lymph nodes underwent activation and clonal expansion within the tumour, prior to PD1 and Lag3 expression, while tumour-associated myeloid cells promoted the formation of a suppressive niche through cytokine secretion and inhibitory T cell interactions. We identified three temporally distinct cancer-associated fibroblast (CAF) populations displaying unique signatures, and verified these in human datasets. In early tumours, immune CXCL12/CSF1 and complement-expressing CAFs supported recruitment of macrophages, whereas contractile CAFs became more prevalent in later tumours. This study highlights the complex interplay and increasing diversity among cells that co-evolve with the tumour, indicating that from early stages of development, stromal cells acquire the capacity to modulate the immune landscape towards suppression.

## Introduction

To aid their growth and development, malignant cells cultivate a supporting niche of ‘normal’ cells, known as the tumour stroma. This niche comprises non-immune cells such as fibroblasts, blood and lymphatic endothelial cells, as well as numerous immune populations^1^. In particular, the balance of anti- vs. pro-tumour leukocytes can dictate tumour fate^2,3^, andin many cases suppressive populations persist to support immune escape and prevent tumour clearance. While, immunotherapies such as anti-CTLA4, anti-PD1 and anti-PD-L1 show efficacy in a large number of melanoma patients, a significant proportion do not respond to this treatment^4–7^. Thus, there remains a critical need to uncover novel therapeutic targets. The numerous mechanisms through which stromal cells promote tumour growth, represent a wealth of opportunities for therapeutic intervention. However, the evolving tumour microenvironment is extremely dynamic, continually adapting to both soluble and mechanical cues which induce significant heterogeneity within the stromal compartment^8^.

In particular, extensive heterogeneity has been reported within tumour fibroblast populations. Cancer associated fibroblasts (CAFs) are the most abundant non-immune stromal component, secreting growth factors, promoting angiogenesis, facilitating metastasis and regulating immune infiltrates^9–15^. Although they express typical fibroblast markers, such as fibroblast activation protein (FAP), platelet derived growth factor receptors α (PDGFRα) and β (PDGFRβ), podoplanin (PDPN), Thy-1 and α-smooth muscle actin (αSMA), no single marker universally identifies all CAFs within the tumour stroma^16–18^. To date, many studies rely on positive selection approaches, in which one or two markers are used to isolate CAFs for functional characterisation. Consequently, these findings likely reflect a sub-population of cells and may bias our perceptions of CAF function. It remains unclear whether fibroblast subpopulations with distinct roles are present in the tumour microenvironment.

Current approaches lack the resolution to visualise the true extent of stromal heterogeneity and may mask rare populations, or cellular phenotypes, that could be critical for tumour survival. Therefore, we have employed single-cell RNA sequencing (scRNAseq) to profile both immune and non-immune stromal populations from the B16-F10 murine model of melanoma. Furthermore, cells were isolated from both primary tumours and draining lymph nodes, at different stages of tumour development, enabling a systems level interrogation of the melanoma microenvironment in real-time. Here, we identified the presence of a diverse immune landscape, in which the composition and phenotype of leukocytes change as the tumour evolves. In particular, effector T cells displaying signs of dysfunction, were detected predominantly in late stage tumours. This work also highlighted significant heterogeneity within the CAF compartment of the primary tumour. Three distinct CAF populations were identified; immune, desmoplastic and contractile, each displaying unique functional and temporal characteristics key to the tumour. At early time points, the ‘immune’’ and ‘desmoplastic’ populations dominated, yet at later stages, the third ‘contractile’ subset became more prevalent. Using a unique database of known ligand-receptor interactions, we investigated communication between different stromal populations to reveal complex interplay between the ‘immune’ CAF subset, macrophages and T-cells, which ultimately contributed to T-cell dysfunction.

## Results

### Identification of stromal populations within the developing tumour microenvironment

Specific immune populations were enriched for, based on surface marker expression, and index sorted from tumours and lymph nodes at day 5, 8 and 11. Isolated single cells were then profiled using Smart-seq2 (Fig. 1a and Extended Data Fig. 1a). To avoid the biases associated with isolation of non-immune stroma, B16-F10 melanoma cells were injected into CAG-EGFP mice exhibiting widespread eGFP expression. This enabled a negative selection approach which did not rely upon expression of surface markers. Tumour and immune cells were removed by selecting GFP^+^ CD45^−^ cells only, with the remaining stromal cells separated into two fractions, based on CD31 expression (Fig. 1a). CD31^+^ cells represented both blood and lymphatic endothelial cells, whereas CD31^−^ cells were largely composed of fibroblasts. Graph-based clustering^19^ of more than 4600 cells that passed the computational quality control (see Methods and Extended Data Fig. 1b), confirmed that populations clustered together with the exception of endothelial cells from tumour and lymph node (Fig. 1b, c and Extended Data Fig. 2). Within each annotated cluster, we could also identify site-determined groupings, specific to either the tumour or lymph node derived cells (Fig. 1b). In particular T cells and dendritic cells exhibited differential clustering between tumour sites and lymph nodes. Moreover, in contrast to recent studies, our approach of sampling over multiple times points, at each site, enabled us to investigate temporal changes between sites and adaptation within each population (Fig. 1b).

**Figure 1.**
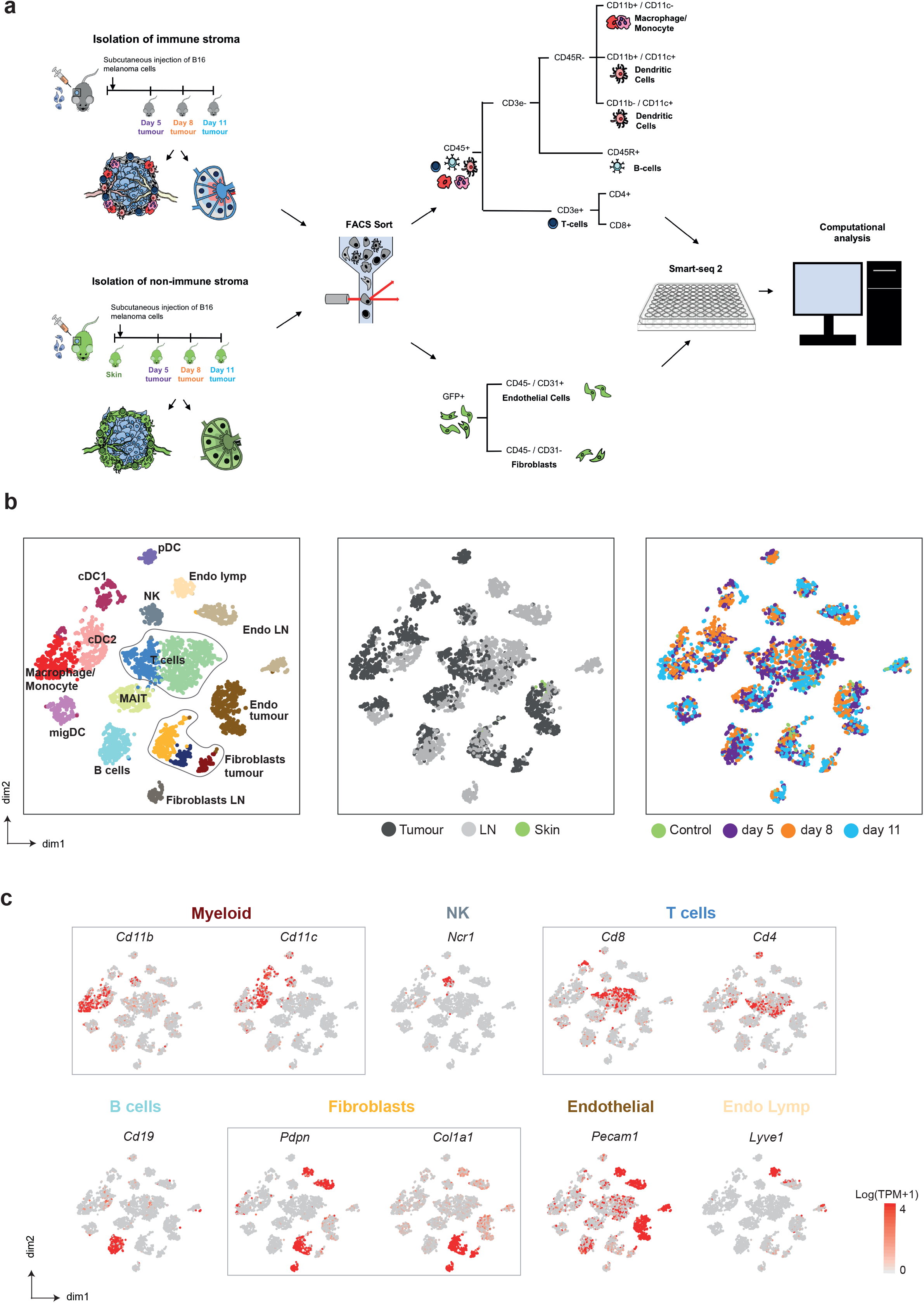
Distinction of melanoma stromal populations with single-cell RNA-seq. **a**, Overview of experimental and sequencing workflow. **b**, tSNE visualisation of all cells sequenced with each cell colour coded for; cell type (left), site of origin (middle), time (right). **c**, Expression of marker genes for each cell type. NK, natural killer; migDC, migratory DC; DC LN, lymph node dendritic cell, cDC1/2, conventional dendritic cell; pDC, plasmacytoid DC; MAIT, Mucosal-associated invariant T cell; Endo lymph, lymphatic endothelial cell; Endo tumour, tumour endothelial cells; endo LN, lymph node endothelium; fibroblast LN, lymph node fibroblast.

### Dynamics of immune stroma

We first sought to delineate relationships within the specific innate immune populations isolated from tumour-associated tissues. Clusters corresponding to macrophages/monocytes, natural killer cells (NK), plasmacytoid DCs (pDC) and conventional DCs (cDCs), were identified based on known markers (Macrophages/Monocytes, *Adgre1*(F4-80); *FcyR1*, NK *Ncr1*; pDCs, *Bst2, Siglech*; cDCs, *Itgax* (*Cd11c*); Fig. 2a, b, c and Supplementary Table 1). Moreover, multiple DC populations were observed that reflect the *Cd11c*+ cDC1 and *Cd11c*+*Cd11b*+ (*Itgam*) cDC2 phenotypes. cDC1 and cDC2 titles were assigned based on expression of known markers including *Clec9a*, *Baft3* (cDC1), *Cd11b* and *Sirpa* (cDC2) (Fig. 2c,b). Two further clusters that lacked lineage markers for adaptive immune cells, as well as ILCs, yet express low *Cd11b* and *Cd11c*, were termed migratory DCs due to expression of the DC transcription factor *Baft3* and upregulation of *Ccr7*.

**Figure 2.**
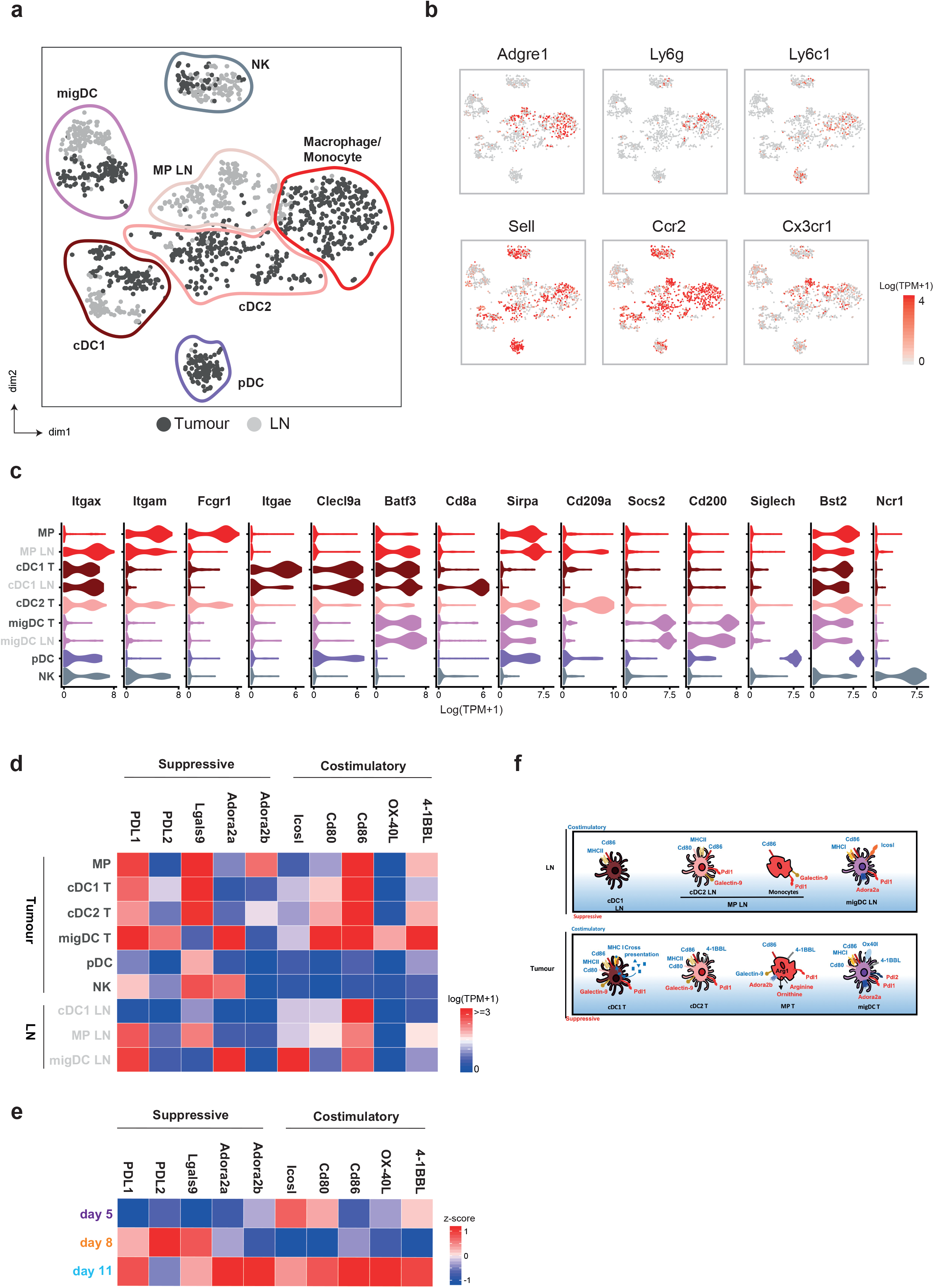
Myeloid cell clusters in the tumour exhibit suppressive characteristics. **a**, tSNE plot of individual myeloid cells colored by site (tumour, dark grey; lymph node, light grey) and clusters marked by coloured lines. **b**, Violin plots showing expression of selected surface marker genes within each cell cluster displayed as Log (TPM+1). **c**, tSNE plots showing expression of selected marker genes for macrophages and inflammatory and resident monocytes. **d**, Heatmap showing mean expression (Log(TPM+1)) of co-stimulatory and suppressive genes for the identified cell clusters. **e**, Heatmap showing relative expression (z-score) of co-stimulatory and suppressive genes in all innate immune cells over time. **f**, Schematic diagram of the costimulatory and inhibitory receptors/ligands expressed on distinct myeloid subpopulations. migDC, migratory DC; DC LN, lymph node dendritic cell, cDC1/2, conventional dendritic cell; pDC, plasmacytoid DC; MP, mononuclear phagocyte.

Each DC population further separated according to their location in either the tumour or draining lymph node (Fig. 2a). Consistent with the steady state, tumour cDC1 cells expressed the dermal DC marker *Cd103* (*Itgae*), whereas the LN population express *Cd8a*, a marker of lymph node resident cDC1 populations. *Cd11b*+ MPs in the LN consisted of *Adgre1*+, *Ccr2*+ macrophages, as well as a *Cd11c*+ resident cDC2 population (Fig. 2b). Investigation into transcriptional phenotypes of myeloid cells revealed that cells located in the tumour, compared to the lymph node were more activated, yet also displayed immunosuppressive properties. Within tumour MPs, no clear delineation between an M1 or the pro-tumour M2 phenotype was observed, yet chemokines involved in T cell recruitment and suppressive mediators such as *Cd274* (PDL1), *Arg1*, *Ido1*, *Adora2a/b* were expressed. Furthermore, while tumour DC populations displayed increased expression and a wider variety of co-stimulatory molecules, than their lymph node counterparts, they lack expression of cytokines required to induce durable T cell responses. This is particularly relevant in regard to cDC1 cells, which can cross-present tumour antigen to cytotoxic T lymphocytes. Indeed, while both tumour and LN cDC1 cells upregulate genes involved in cross-presentation pathway, including components of the proteasome (*Tap1*, *Tap* and *Sec61*), expression of these genes was higher in the tumour subset (Extended Data Fig. 3d). Although this may indicate that tumour cDC1 populations have increased potential to activate anti-tumour lymphocytes, tumour DCs also expressed immunosuppressive molecules such as *Cd274* (PDL1), *Pdcd1lg2* (PDL2) and *Lgals9* (galectin-9), known to induce T cell exhaustion (Fig. 2d). Interestingly, across all myeloid populations, expression of immunosuppressive molecules increased at later time points, whereas co-stimulatory molecules were expressed consistently throughout tumour development. This indicates that tumour resident myeloid populations are present and activated at early stages of tumour growth, yet become more suppressive as the tumour progresses.

T cell populations from tumours and draining lymph nodes were transcriptionally distinct, clustering based upon their subtype, but also location (Fig. 3a). At the lymph node, T cells exhibited a more naive phenotype compared to those present at the primary tumour (Fig. 3b). While tumour resident CD4+ T cells were more activated, a significant proportion highly expressed Treg-associated genes at the tumour (Fig. 3b). Similarly, within the CD8+ T cell compartment, those at the tumour were also more activated, expressing high levels of *Ifng* (IFNγ), *Prf1* (perforin) and *Gzmb* (granzyme B). However, these cells were also less functional, as evident by expression of *Pdcd1* (PD1), *Lag3* and *Tim3* (Fig. 3b). To identify transcriptional adaptations in CD8+ T cells, at different stages of tumour development, we performed pseudotime analysis that revealed a trajectory of gene expression associated with functional changes in these cells. This confirmed that the majority of T cells within the lymph node were naive displaying high expression of *Sell* and *Tcf7* (Fig. 3c and d). Arrival at the tumour corresponded with acquisition of activation signatures, including upregulation of *Ifng* (IFNγ) and *Gzmb* (Granzyme B). Furthermore, T cell receptor sequence analysis identified clonal expansion (Fig. 3c and d) specifically within tumours at later time points. This was accompanied by expression of the proliferation marker *Mki67* and exhaustion markers *Pdcd1*, *Lag3* and *Tim3* (Fig. 3c and d). Interestingly, a subset of the potentially exhausted CD8^+^ T cells also showed expression of *Entpd1* (CD39), which was recently identified as a marker to distinguish tumour-specific and bystander CD8^+^ T cells ^20^. Together, these results indicate that T cell recruitment from the LN is followed by activation and subsequent functional defects *in situ*. These functional defects correspond with the gain of immunosuppressive properties in myeloid populations at later time points, indicating that the immune stroma transitions from immunogenic to suppressive phenotypes.

**Figure 3.**
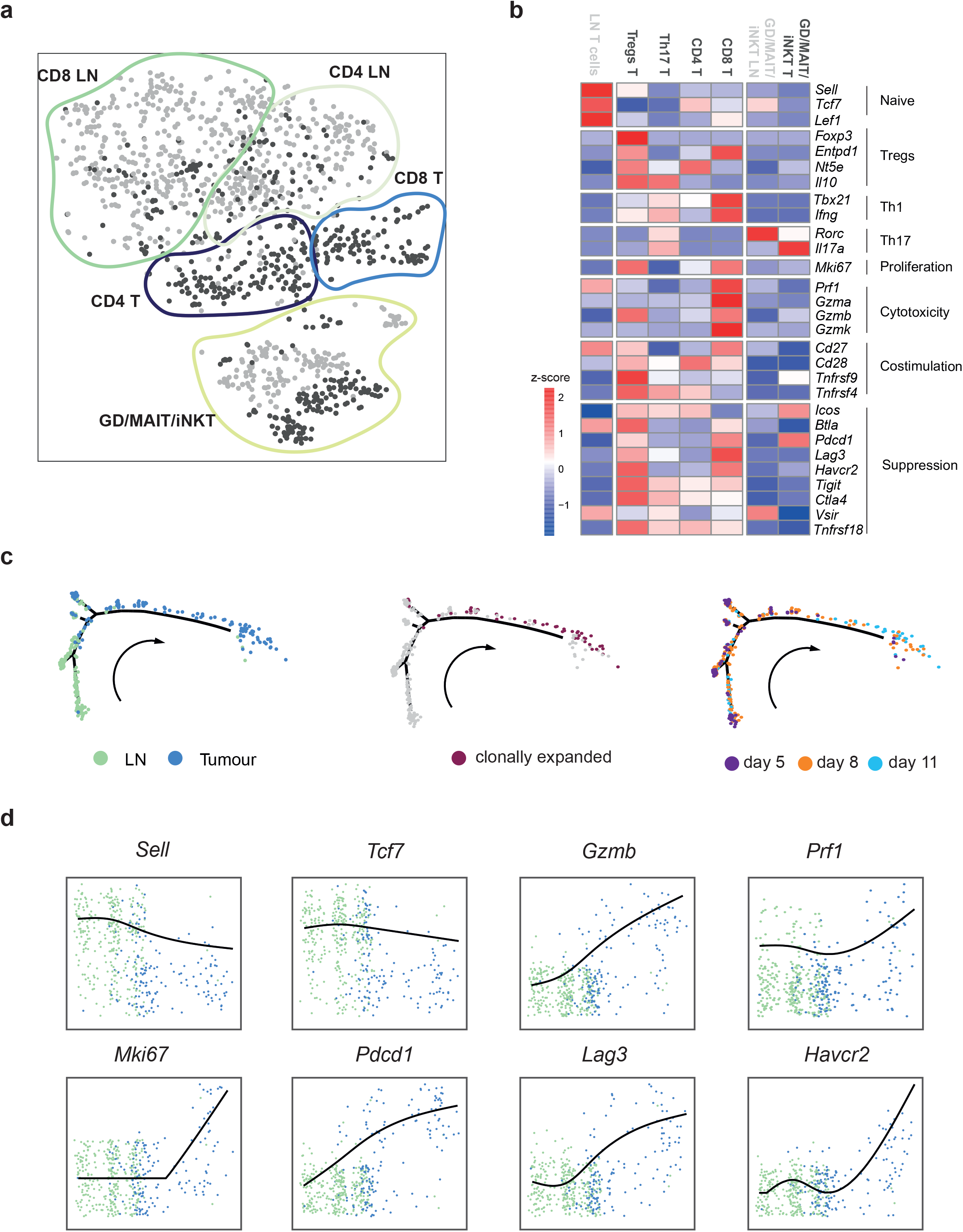
T cells recruited from lymph nodes are activated *in situ*. **a**, tSNE plot of individual T cells colored by site (tumour, dark grey; lymph node, light grey) and annotated subpopulations marked by coloured lines. **b**, Heatmap showing relative expression (z-score) of functional gene groups for cell clusters. **c**, Pseudotime analysis of CD8+ T cell gene trajectories coloured by site (left), clonal expansion (middle) and tumour stage (days, right), arrow indicates time direction. **d**, Expression of activation-associated genes along the inferred pseudotime coloured by site; lymph node (green), Tumour (blue).

### Non-immune stroma comprise three distinct functional populations

As the non-immune stromal components are emerging as immune modulators, we also examined this compartment during tumour progression, focussing on the CAFs. Across all time points, we identified three distinct CAF populations referred to as CAF 1, 2 and 3 (Fig. 4a). As expected, expression of commonly used CAF markers was extremely variable across the fibroblasts (Fig. 4e and Extended Data Fig. 6a), yet, expression of specific marker combinations correlated with individual clusters. CAF1 could be distinguished from CAF3 by its high levels of *Pdpn*, *Pdgfrα* and *Cd34*, while *Acta2* (αSMA) was strongly expressed by the latter population. However, CAF2 represents an intermediate population that was *Pdpn^+^ Pdgfrα^+^* and displayed low expression of *Acta2* and *Cd34* (Fig. 4e).

**Figure 4.**
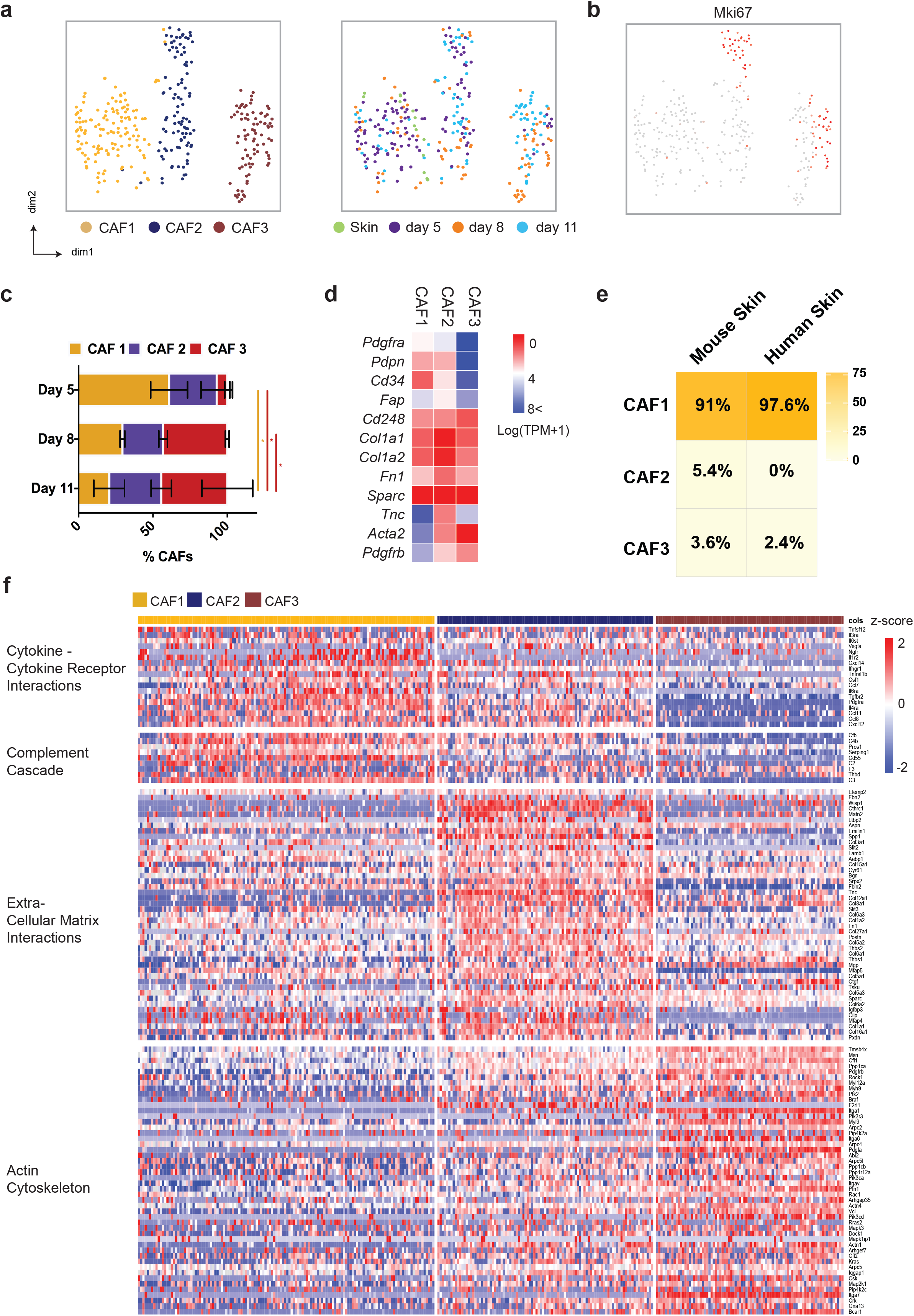
Distinct fibroblast clusters identified in melanoma tumours. **a**, tSNE plot of sequenced fibroblasts from tumours coloured by their associated cluster (left) or by tumour time point (right) **b**, tSNE visualisation of the proliferation marker Mki67 in the CAFs. **c**, bar plot depicting the ratio of CAF populations at each time point examined where the size of each coloured bar is proportional to percentage of total CAFs each population represents. Data presented as mean ± SEM. * P<0.05 (Two way anova with Tukey post-hok test). **d**, Heatmap showing average expression (Log(TPM+1)) of typical CAF markers. **e**, Heatmap depicting logistic regression analysis of normal mouse skin, indicating which of the 3 CAF clusters these cells are most similar. **f**, Heatmap of gene ontology pathways for differentially expressed genes in each cluster; cytokine-chemokine receptor interactions, complement cascade, Extracellular matrix interactions and actin cytoskeleton. Columns represent individual cells, rows display z scores.

Importantly, each cluster displayed distinct functional signatures (Fig. 4f, Extended Data Fig. 5c,d), indicating that fibroblast populations may have specific roles within the tumour microenvironment. CAF 1 (*Pdpn^+^ Pdgfra^+^ Cd34^high^)* upregulated genes involved recruitment and regulation of immune cells, including the cytokines *Cxcl12*, *Csf1* and *Ccl8*, cytokine receptors *Il6ra* and *Il6st*, as well as components of the complement cascade *C3*, *C2 and C4b*. Thus, CAF1 may engage in immune cross-talk. In contrast, CAF2 (*Pdpn^+^ Pdgfra^+^ Cd34^low^)* expressed genes encoding extracellular matrix (ECM) components including numerous collagen family members and *Postn* and *Tnc*. These ECM components are strongly associated with a fibrotic matrix, a feature common to developed tumours ^21^. Thus, it is possible that this CAF population drives the desmoplastic reaction associated with tumour development. CAF3 (*Acta2^high^*) was enriched for genes involved in regulation and rearrangement of the actin cytoskeleton. In particular, this cluster upregulated *Rock1*, *Mlc2* and *Mlck*, which are responsible for the contraction of actin stress fibres. Thus, CAF 3 likely represents a more contractile fibroblast subset. CAF3 also expressed some pericyte-associated markers such as *Cspg4* (Ng2), *Mcam* and *Rgs5* (Extended Data Fig. 7a), leading us to consider whether this population may contain pericytes. The same markers were also observed in the *Pdpn*+ fibroblasts (FRCs) in the lymph nodes however, (Extended Data Fig. 7b) indicating that, and consistent with previous reports, their expression is not limited to pericytes. Thus, to identify whether CAF3 represent a pericyte or fibroblast population, we examined expression of αSMA, Ng2 and Mcam in relation to the endothelial marker CD31. While these were observed surrounding vessels in adjacent skin, they were rarely associated with intratumoural vessels, but could be detected in peritumoural spindle-shaped cells distinct from the vasculature (Extended Data Fig. 7 c-e).

Our approach highlighted the dynamic nature of CAF populations within a developing tumour. Although each CAF population was detected throughout the time course, different clusters dominated at specific time points. Early day 5 tumours were primarily comprised of fibroblasts from the *Pdpn^+^ Pdgfra^+^* CAF1 and 2, whereas the *Acta2^high^* CAF3 population was largely restricted to later stages, implying a selective enrichment in developed tumours. This enrichment may be supported, in part, by our observation of proliferation specifically within CAF2 and 3 (Fig. 4b). Proliferation of a subset of CAFs, within the tumour microenvironment, was confirmed by incorporation of the thymidine analogue, EdU (Extended Data Fig. 6c). Moreover, the majority of both mouse skin and human skin fibroblasts resembled the cells from CAF1 (Fig. 4e). Together, this data illustrates that the CAF compartment and its associated functions are dynamic, adapting to localised cues and the changing needs of an evolving tumour.

To validate the existence of these different populations in the tumour microenvironment, we first confirmed each subset based on their unique marker repertoire. Consistent with sequencing data, confocal imaging revealed that CAF markers PDPN and PDGFRα largely colocalised, while expression of αSMA was more distinct (Fig. 5a). The ‘immune’ CAF1 marker CD34, colocalized with both PDPN and PDGFRα, indicating the presence of a CD34^high^ subpopulation. Furthermore, a distinct CD34^high^αSMA^low^ CAF subset could be clearly distinguished (Fig. 5a). Although at the RNA level, PDPN^+^ PDGFRα^+^ CD34^high^ CAFs are αSMA^−^ (Extended Data Fig. 6a), we observed some colocalization between these four markers at the protein level. This may represent the intermediate PDPN^+^ PDGFRα^+^ CAF2 population, which also expressed low levels of CD34 and αSMA.

**Figure 5.**
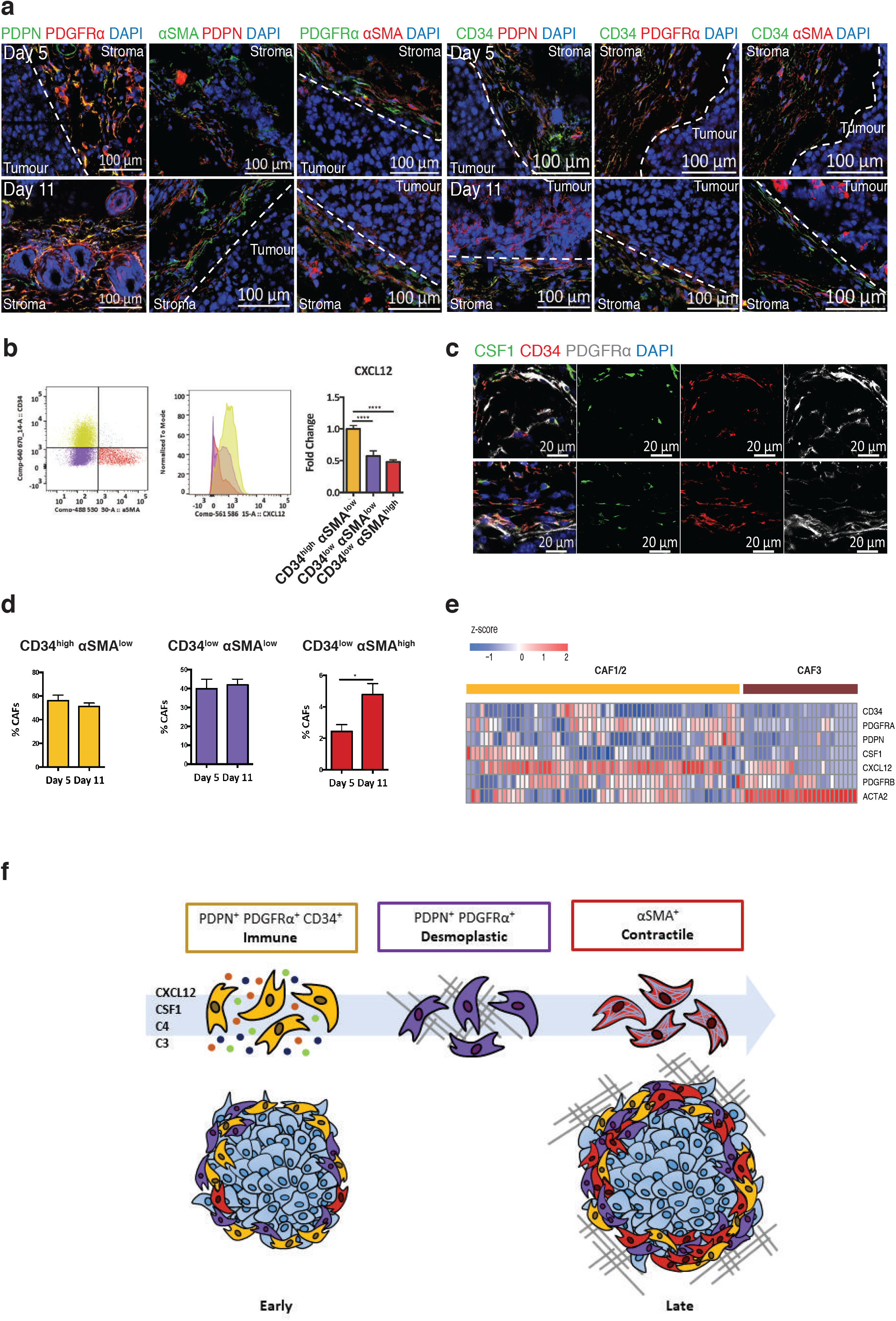
Fibroblast subtypes in murine are mirrored in human disease. **a**, Representative confocal images of PDPN, PDGFRα and αSMA in combination (left panel) or CD34 in combination with either PDPN PDGFRα or αSMA (right panel) in day 5 and day 11 tumours. Dashed line indicates the tumour border. Scale bars 100μm. **b**, Verification of populations by flow cytometry. Representative plots depicting the method by which each population was distinguished, based on CD34 and αSMA expression (left panel) and a histogram showing intracellular CXCL12 expression in each population (middle). Fold change in mean fluorescence of normalised to the CD34^high^ αSMA^low^ population at day 11 (right), is also shown. **c**, Representative confocal image of CSF1 expression in αSMAlow CD34+ CAF populations in day 5 and day 11 tumours. Scale bars 20um. **d**, Flow cytometry quantification of the proportion of each CAF population at day 5 and day 11 tumours, displayed as as the percentage of the total CAF population. **e**, Heatmap displays expression (z scores, blue to red) of key markers and cytokines across CAFs clusters identified in human melanoma. **f**, Schematic diagram of the three CAF subpopulations. (a, c, d representative image from at least n=3 independent mice). (b) n=12 independent mice. (d) Day 5: n=9 tumours from 8 independent mice, Day 11: n=11 independent mice. Data presented as mean ± SEM. * P<0.05 as calculated using either a one way anova with Dunnett’s post-hok test (b) or Student’s T-test (d)

To examine the inflammatory phenotype associated with the CAF1 population in more detail, we focused on two highly expressed immune modulatory factors, CXCL12 and CSF1. Using flow cytometry we identified each population using their differential expression of CD34 and αSMA. Tumour fibroblasts were identified using multiple CAF markers, after exclusion of other stromal populations (Extended Data Fig. 8a), and divided into CD34^high^ αSMA^low^ (CAF1), CD34^low^ αSMA^low^ (CAF2) and CD34^low^ αSMA^high^ (CAF3) subsets. Reflecting our sequencing data, this confirmed CXCL12 expression was highest in the CAF1 subset, followed by intermediate expression in CAF2 and low expression in CAF3 (Fig. 5b), this was further verified with RNAscope showing localization of CXCL12 and CD34^+^ at the RNA level in situ (Extended Data Fig. 6d). Confocal imaging also confirmed CD34^+^ CAFs as a source of CSF1 in the tumour stroma both at the protein (Fig. 5c) and RNA levels (Extended Data Fig. 6d). Having validated the presence of these functionally distinct populations, we next evaluated their prevalence at different stages of tumour development (Fig. 5e). This showed that, as a percentage of the total CAF population, the proportion of of CD34^low^ αSMA^high^ CAF3 subset was greater at day 11 compared to day 5. This data supports our earlier proposal of a dynamic fibroblast niche, which evolves alongside its malignant tumour core.

Together, these data have identified the presence of distinct CAF populations that dynamically co-evolve with the tumour to support its changing requirements (Fig. 5g) and indicate that CAFs acquire the capacity to influence the tumour immune landscape from early stages of development.

### Cross-talk between ‘immune’CAFs in infiltrating myeloid cells

Next, we sought to elucidate the potential functional consequences of specific stromal populations to the ensuing immune response. Thus we focused on the early CAF1 “immune” population and examined the cross-talk with responsive immune populations recruited to the tumour. To systematically study interactions within the tumour microenvironment, we predicted cell-cell communication networks based on CellPhoneDB, a manually curated repository of ligands, receptors and their interactions integrated with a statistical framework to infer cell-cell communication networks from single cell transcriptomic data (Vento-Tormo, Efremova et al, Nature, in press). This approach highlighted likely interactions involved in angiogenesis, immune cell recruitment and immune modulation between stromal populations in the tumour (Fig. 6a, Supplementary Table 4).

**Figure 6.**
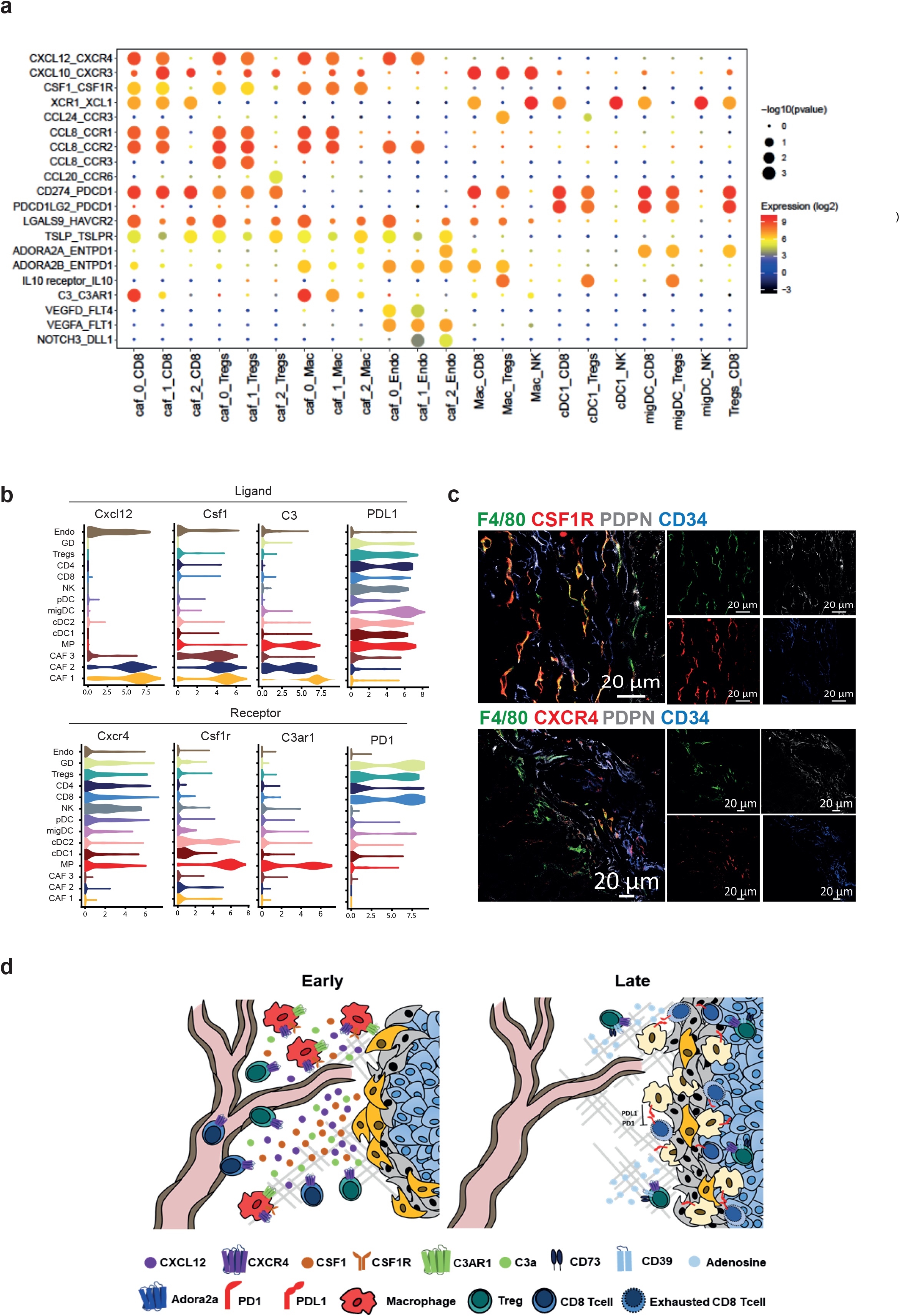
Stromal crosstalk supports the development of an immune suppressive niche. **a**, Overview of selected statistically significant specific interactions between stromal subsets and other cell types using a cell-cell communication pipeline based on CellPhoneDB. Size indicates *p*-values and colour indicates the means of the receptor/ligand pairs between two clusters. **b**, Violin plots displaying expression of ligands *Cxcl12*, *Csf1, C3* and PDL1 and cognate receptors *Cxcr4*, *Csf1r, C3ar1* and PD1 on respective stromal populations. **c**, Confocal images of representative tumour-tissue borders. CSFR1 or CXCR4 expressing macrophages located proximally to CD34+ CAFs (green, F4/80; red, CXCR4 or CSF1R; white, podoplanin; blue CD34. Scale bars, 20um. **d**, Schematic diagram of the dynamic cross-talk identified within the tumour microenvironment.

We identified CAF-immune interactions, for example between C3/CXCL12/CSF1-expressing CAFs enriched in early stages of tumour development, and macrophages positive for CXCR4, CSFR1, C3AR1 respectively (Fig. 6a, b). IF imaging of tumour sections allowed us to correlate single cell data with location in situ. Indeed, at the protein level, both CSFR1^+^ and CXCR4^+^ macrophages were detected in close contact with CD34^high^ fibroblasts in the tumour stroma (Fig. 6c). The combination of transcriptome profiling and cell-cell communication pipeline enabled us to assign these immune interactions specifically to the CAF1/2 subpopulations. Further chemokine-receptor interacting pairs, identified as statistically significant, occured between the immune CAF1 subpopulation, myeloid, Treg and CD8^+^ T cells (Fig. 6a). The recruited macrophages exhibited the capacity to both attract T cells, via specific cytokine-receptor signals such as CXCL10, and suppress their function through the PDL1-PD1 axis (Fig. 6a and d). Our approach highlighted additional cell-cell communications enriched between tumour infiltrating immune such as recruitment of NK cells through cDC1-cell derived chemokines receptors XCR1^22^. Additionally, we found that the Tregs express high levels of *Nt5e* (CD73) and *Entpd1* (CD39, Fig. 6a and Fig. 3b), which act together to convert ATP to adenosine who release has been shown to dampen the immune system^23^. The adenosine receptors *Adora2a* and *Adora2b* were found upregulated on the migratory DCs and the macrophages respectively.

Collectively, these findings provide new insights into the complex interplay among cells within the evolving tumour microenvironment (Fig. 6d), where multiple immunosuppressive mechanisms coexist within an increasingly heterogeneous stromal compartment.

## Discussion

It is becoming increasingly evident that non-malignant stromal cells such as endothelial cells, fibroblasts, and infiltrating immune cells found within a tumour, provide significant and varied supporting roles as disease progresses. The heterogeneity and dynamic nature of the tumour microenvironment can make identification of the roles of the different stromal components challenging. The emergence of scRNA-seq has enabled new insights into tumour biology not detectable by previous methods, and has been key to reveal the true degree intratumoural heterogeneity ^24,25^.

In this study, we used a single-cell transcriptomic approach to characterise the stromal landscape within the evolving microenvironment. Our scRNA-seq analysis revealed the gradual development of a suppressive immune microenvironment and defined fibroblast subsets with distinct functional signatures. This approach also highlighted the complexity of cross-talk between the different stromal components as a tumour evolves.

Both clinical studies and the success of immune checkpoint inhibitors have emphasised the importance of the immune system, particularly T cells and macrophages, in deciding tumour fate or response to therapy. While clinical studies have repeatedly demonstrated the presence of exhausted T cells with poor prognosis ^2,3^, the steps leading to this point and sites of activation are less clear. Here we showed distinct gene profiles between sites, with lymph nodes acting as a source of naive T cells. Once at the tumour, pseudotime analysis illustrated the trajectory of T cell development within the evolving tumour microenvironment from naive, through clonal expansion and activation (enriched granzyme and IFN expression) phases, to upregulation of exhaustion markers in late tumours (PD1 and Lag3). A diverse repertoires of myeloid cells were observed within the tumour, and similar to T cells, tumour myeloid populations were more activated than in the lymph node displaying high levels of phagocytosis, antigen presenting and co-stimulatory associated genes. Once at the tumour however, an increase in the level of suppressive factors produced, likely in response to local cues, was detected. Many of these cues served to act upon infiltrating T cell populations to confound the suppressive environment already developing. Moreover, our dynamics data indicates that inhibitory signalling commenced in later phases of tumour development, coinciding with the emergence of T cell dysfunction.

While infiltrating immune populations can have a profound effect on tumour fate, a growing body of evidence indicates that CAFs play a supporting role in the tumour microenvironment^26^. To date, few studies have investigated whether fibroblast phenotypes change as the tumour develops. Our analysis revealed the existence of three CAF subsets which possess unique characteristics and temporal dynamics, indicative of distinct, specialized roles. Based on their transcriptional signatures, these populations were termed “immune”, “desmoplastic” and “contractile”. The ‘immune’ CAF1 population, which was detected from early stages of tumour development, upregulated cytokines CSF1 and CXCL12, as well as complement components C3 and C4b, which are known to recruit and regulate immune cells. Conversely, the desmoplastic population upregulated ECM components and may be responsible for the production of a fibrotic matrix. Finally, the third population, dominated at later stages of tumour development in well-established lesions, expressed genes involved in the contraction of actin stress fibres, indicating a contractile phenotype.

It is likely that the alteration in population dynamics is induced by concomitant changes in the developing tumour. New environmental factors, such as nutrient availability and hypoxia arise, transforming the secretion profile of malignant cells. Furthermore, changes in the phenotype and function of the immune stroma, may add to the local cytokine milieu. Biophysical cues also likely contribute to adaptation of CAF populations. It has been reported that matrix rigidity is critical for the maintenance of CAF phenotypes ^34^, and that mechanical and soluble cues are required for the induction of αSMA expression ^35,36^. Thus it is likely that the combination of a stiff matrix, produced by ‘desmoplastic’ CAFs, and cytokine exposure, may upregulate αSMA expression and expansion of ‘contractile’ CAFs in later tumours. No single commonly applied marker identified all CAF subsets. Instead the 3 populations were distinguished based on their discrete expression of CAF marker combinations. While both ‘immune’ and ‘desmoplastic’ CAFs express PDPN and PDGFRα, the ‘immune’ population was distinguished based on its high expression of CD34. The ‘contractile’ population is largely negative for these markers and instead expresses αSMA.

Interestingly, and consistent with other studies, this population also shared some common markers with pericytes, such as Ng2, Mcam and Rgs5^37^. While this could imply pericyte contamination, CAF3 also produced matrix components such as *Col1a1, Col1a2* (*collagen1) Fn1 (fibronectin1)* and *Sparc*. Although expression of these genes was lower in CAF3 compared with the desmoplastic CAF2 population, production of these proteins is associated with a fibroblast phenotype. Pericytes are located on the surface of blood vessels where they provide structural support, as well as regulating endothelial cell phenotypes. Like fibroblasts, these cells represent a heterogeneous population that is difficult to distinguish by a specific marker. While markers such as NG2 and RGS5 are often used to identify pericytes, their expression is context-dependent, varying between tissues and during pathology ^38^. Furthermore, many markers, such as αSMA, PDGFRβ, Ng2 and Mcam are shared by both pericytes and activated myofibroblasts, making it difficult to distinguish one from the other ^39,40^. This was reflected in our data, in which expression of multiple pericyte markers was observed in the CAF3 population as well as in PDPN+ lymph node FRCs. A similar phenomenon was reported in a subcutaneous model of breast cancer, in which expression of pericyte markers was also observed in FRC populations ^41^. Thus, the most robust method to differentiate between these cell types is to assess whether they are associated with endothelial vessels. In our melanoma model, we observed aSMA, NG2 and MCAM positive cells both associated with vessels and in more peripheral locations. However, the number of vessel associated pericytes was very small, as the majority of vessels were composed of endothelial cells alone. Consequently, CAF3 may embody a mixed population of mesenchymal cells, containing both pericytes and fibroblasts, that share similar surface marker expression and functional properties. Interestingly, the close relationship between these two cell types has led to the suggestion that pericytes may differentiate into activated myofibroblasts during pathology. In both liver and kidney fibrosis, as well as in tumour models, upon the initiation of fibrosis or during growth of malignant cells, pericytes dissociate from the endothelium and begin to express the markers such as αSMA and produce collagen ^42–45^. Therefore, it is possible that fibroblasts within CAF3 may arise from pericyte origins. Overall, this data highlights the limitations of using single marker approaches to isolate and characterise mesenchymal cells, which can lead to contamination and selection bias.

Importantly, and relevant to the clinic, our murine data was mirrored in the setting of human melanoma, with shared patterns of CAF marker expression ^24^. Here, CAFs expressing PDPN, PDGFRα and CD34 also clustered together, whereas those expressing αSMA were more distinct. Furthermore, these PDPN^+^PDGFRα^+^CD34^+^ CAFs displayed high expression of CXCL12. Other immunomodulators such as PD-1 ligands and, in particular, complement components were observed in both systems suggesting that the immune function of these populations are a conserved feature and retained in human melanoma. Although the CAF 1/2 clusters were not as distinct as the mouse model, the cohort of CAFs was much smaller. This discrepancy can be explained by the fact that many of these patients had received immunotherapies prior to resection ^24^, and in light of the immunomodulatory capacity of these clusters, we cannot rule out an effect of treatment on the wider stromal landscape.

Our findings also compliment recent investigations into CAF heterogeneity in a range of different solid cancers, each reporting a distinct αSMA^+^ fibroblast phenotype ^25,46,47^. Moreover, αSMA^−^ fibroblasts highly expressing ECM components, similar to our ‘desmoplastic’ population have been described in both colorectal and head and neck cancers ^25,46^. Furthermore, in a preclinical model of pancreatic ductal adenocarcinoma (PDAC), another αSMA^−^ CAF population was shown to display an inflammatory profile ^47^. Similarly, a subpopulation of CXCL12-secreting fibroblasts was reported in human breast cancer. However, in contrast to our results, a proportion of this CAF population was αSMA+ ^48^. This suggests that the three populations we have identified may be a universal feature of the tumour stroma in a variety of cancer types. Subtle difference between the fibroblast populations reported, such as marker expression and the range of cytokines produced, likely reflect the local milieu of soluble and mechanical cues, as well as environmental pressures unique to the tumour type.

Sequencing of paired immune and non-immune stroma provided the opportunity to investigate signalling between different stromal compartments. Using a recently reported database of receptor-ligand interactions, we were able to infer cross-talk between the ‘immune’ CAF subset and *Cd11b*^+^ *Cd11c*^−^ macrophages, via the CXCL12-CXCR4, CSF1-CSF1R and C3-C3ar1 axes. While both of CXCL12 and CSF1 are reported to recruit macrophages to the tumour stroma and induce a suppressive phenotype, the role of the complement cascade in the tumour microenvironment is more ambiguous ^29,49–52^. Complement components are typically thought to aid immune clearance by increasing opsonization and phagocytosis, formation of the membrane attack complex and recruitment of multiple immune populations ^53^. However, in a malignant context, the complement cascade has been shown to promote tumorigenesis and induce immune suppression. In particular, complement components C3a and C5a and their cognate receptors C3AR1 and C5AR1 are linked with recruitment of suppressive myeloid cells and T cell dysfunction ^54–58^. Interestingly, the CAF1 subset represented the greatest source of C3 in the primary tumour. This component acts upstream within the complement cascade, inducing the cleavage and activation of downstream factors. Thus, while CAF1 secreted C3 may recruit macrophages to the tumour stroma, C3 activation of complement components, such as C5a, may broadly suppress immune function.

Thus, ‘immune’ CAFs may aid in the recruitment and polarisation of suppressive macrophage populations, which then further contribute to the development of immune privilege by suppressing T cell function. Furthermore, the CAF1 subpopulation may directly induce T cell dysfunction by activation of the complement cascade. This additional layer of immune regulation by CAFs is also consistent with recent studies in pre-clinical models of PDAC and melanoma, which suggested that blockade of either CXCL12-CCR4, CSF1-CSF1R or complement receptors, act to synergise with checkpoint inhibitors ^57^,32,59. With inhibitors targeting both CXCR4 and CSF1R, as well as numerous components of the complement cascade, currently in clinical trials (for example, CXCR4: AMD3100 and PF-06747143, CSF1R: JNJ-40346527 and PLX5622, C3: APL-1 APL-2), enrichment of ‘immune’ CAFs may highlight patients that would benefit from this treatment, in combination with checkpoint immunotherapies.

In summary, we have demonstrated the power of scRNAseq to define the tumour stromal landscape, highlighting the dynamic and adaptive nature of both immune and non-immune stroma within an evolving tumour microenvironment, and revealedpotential cross-talk between these two compartments. We identified 3 CAF clusters with distinct functional and temporal features; the immune subset supporting recruitment and induction of an immunosuppressive macrophage phenotype providing an alternative, indirect mechanism to dampen T cell mediated anti-tumour immunity.

## Materials and Methods

### Mouse models

Animals were housed in accordance with UK regulations and experiments were performed under project licences PPL 80/2574 or PPL P8837835. The C57BL/6 derived B16.F10 melanoma cell line was purchased from American Type Culture Collection (ATCC) and cultured in Dulbecco’s Modified Eagle medium (DMEM, Life Technologies), supplemented with 1% Penstrep and 10% FBS. 2.5 ×10^5^ B16 cells were injected, subcutaneously, into the shoulders of either wild type (WT) C57BL/6 mice, or C57BL/6-Tg(CAGEGFP)131Osb/LeySopJ mice (Jackson Laboratory). After 5, 8 and 11 days animals were sacrificed and tissues collected for analysis. In addition, skin was also taken from nontumour bearing mice.

### Tissue Processing

Tumours were mechanically dissociated and digested in 1mg/ml collagenase D (Roche), 1mg/ml collagenase A (Roche) and 0.4mg/ml DNase (Roche) in PBS, at 37°C for 2hs. Lymph nodes were mechanically dissociated and digested with 1mg/ml collagenase A (Roche) and 0.4mg/ml DNase (Roche) in PBS, at 37°C. After 30 mins, Collagenase D (Roche) was added (final concentration of 1mg/ml) to lymph node samples and digestion was continued for a further 30 mins. EDTA was added to all samples to neutralise collagenase activity (final concentration (5mM) and digested tissues were passed through 70μm filters (Flacon) ready for staining.

### Isolation of Single Cells

Single cells were isolated from processed tissues using fluorescence-activated cell sorting (FACS). Once processed, samples were incubated with a fixable fluorescent viability stain (Life Technologies) for 20mins (diluted 1:1000 in PBS) prior to incubation with conjugated primary antibodies for 30 mins at 4°C. Antibodies were diluted in PBS 0.5% BSA according to table SX. Stained samples were index sorted, using the BD influx flow cytometer system, Single-cells were sorted in 2μl of Lysis Buffer (1:20 solution of RNase Inhibitor (Clontech, cat. no. 2313A) in 0.2% v/v Triton X-100 (Sigma-Aldrich, cat. no. T9284)) in 96 well plates, spun down and immediately frozen at −80 degrees.

### Preparation of cDNA and sequencing

Reverse transcription and cDNA pre-amplification were performed according to the SmartSeq2 protocol ^60^ to obtain mRNA libraries from single-cells. Oligo-dT primer, dNTPs (ThermoFisher, cat. no. 10319879) and an ERCC RNA Spike-In Mix (1:50,000,000 final dilution, Ambion, cat. no. 4456740) were then added. Reverse Transcription and PCR were performed as previously published ^60^, using 50U of SMARTScribe™ Reverse Transcriptase (Clontech, cat. no. 639538). cDNA libraries were prepared using the Nextera XT DNA Sample Preparation Kit (Illumina, cat. no. FC-131-1096), according to the protocol supplied by Fluidigm (PN 100-5950 B1). Single cell libraries were pooled, purified using AMPure XP beads (Beckman Coulter) and sequenced on an Illumina HiSeq 2500 aiming for and average depth of 1 Million reads/cell (paired-end 100-bp reads).

### Single-cell RNA sequencing analysis

The SmartSeq2 data was quantified with Salmon^61^ (version 0.8.2), using the GENCODE mouse protein-coding transcript sequences. Transcript Per Million (TPM) values reported by Salmon were used for the quality control of the samples. In order to get the endogenous TPM values, we removed the ERCC’s from the expression table and scaled the TPM’s so that they sum to a million. Cells with less than 1500 detected genes and for which the total mitochondrial expression exceeded 20% were excluded from further analysis. Genes that were expressed in less than 3 cells were also removed.

Downstream analysis such as, clustering based on SNN graph-based clustering, differential expression analysis and visualisation were performed using the Seurat package^19^ (version 2.3.4) implemented in R. Clusters were identified using the community identification algorithm as implemented in the Seurat "FindClusters" function. Differential expression analysis was performed based on the Wilcoxon rank sum test. Clusters were annotated using canonical cell type markers. Two clusters of dDC2 in the tumour represented the same cell type and were therefore merged.

Trajectory modelling and pseudotemporal ordering of cells was performed with the Monocle 2 R-package^62^ (version 2.8.0). The most highly variable genes were used for ordering the cells. Potential doublets and contaminating melanocytes and keratinocytes were excluded.

We also removed a cluster for which the top markers were genes associated with dissociation-induced effects.

To further identify subpopulations, we reanalysed the T cells, innate immune cells (myeloid and NK) and the CAFs separately, using the same workflow as described above. To account for the cell cycle heterogeneity in the T cell subsets. a cell cycle score was calculated for each cell and this score was then regressed out. We used the function “AddModuleScore” from Seurat and the list of G2M associated genes from Scaldoen et al. to calculate a cell cycle score for each cell.

Gene Set Enrichment Analysis (GSEA) (software.broadinstitute.org/gsea/index.jsp) was performed on genes that were differentially expressed between clusters, with a p value < 0.05. Overlap with canonical GO categories (CP:BIOCARTA, CP:KEGG, CP:REACTOME) was assessed and the False Discovery Rate (FDR) calculated.

### T-cell receptor (TCR) analysis

The TCR sequences for each single T cell were assembled using TraCeR^63^ which allowed the reconstruction of the TCRs from scRNA-seq data and their expression abundance (transcripts per million, TPM), as well as identification of the size, diversity and lineage relation of clonal subpopulations. In total, we detected 77 TCR sequences with at least one paired productive αβ or gamma-delta chain. Cells for which more than two recombinants were identified were excluded from further analysis.

### Cell cycle analysis

The pair-based prediction method described by Scialdone *et al^64^*. and implemented in the R package scran was used to assign each cell a cell cycle stage. Briefly, using a training data, pairs of marker genes are identified such that the expression of the first gene in the training data is greater than the second in certain cell cycle stage but less than the second in all other stages. For each cell then, the method calculates the proportion of all marker pairs where the expression of the first gene is greater than the second in the test data.

### Putative interactions between cell types

To enable a systematic analysis of cell-cell communication, we used CellPhoneDB (Vento-Tormo, Efremova et al., Nature, in press). CellPhoneDB is a manual curated repository of ligands, receptors and their interactions, integrated with a new statistical framework for inferring cell-cell communication networks from single cell transcriptome data. Briefly, in order to identify the most relevant interactions between cell types, we looked for the cell-type specific interactions between ligands and receptors. Only receptors and ligands expressed in more than 10% of the cells in the specific cluster were considered. We performed pairwise comparisons between all cell types. First, we randomly permuted the cluster labels of all cells 1000 times and determined the mean of the average receptor expression level of a cluster and the average ligand expression level of the interacting cluster. For each receptor-ligand pair in each pairwise comparison between two cell types, this generated a null distribution. By calculating the proportion of the means which are "as or more extreme" than the actual mean, we obtained a *p*-value for the likelihood of cell type-specificity of a given receptor-ligand complex. We then prioritized interactions that are highly enriched between cell types based on the number of significant pairs and manually selected biologically relevant ones. For the multi-subunit heteromeric complexes, we required that all subunits of the complex are expressed (using a threshold of 10%), and therefore we used the member of the complex with the minimum average expression to perform the random shuffling.

### Mouse skin fibroblasts from healthy mice

Skin samples from two 8-week old female C57BL/6 mice were processed, first by mechanical processing, followed by 2 h incubation with 0.5% collagenase B (Roche; 11088815001). Cells were then counted and loaded on the 10x Chromium machine. Libraries were prepared following the Chromium Single Cell 3′ v2 Reagent Kit Manual^65^. Libraries were sequenced on an Illumina HiSeq 4000 instrument with 26 bp for read 1 and 98 bp for read 2.

Droplet-based sequencing data was aligned, filtered and quantified using the Cell Ranger Single-Cell Software Suite (version 2.2.0), against the mouse reference genome provided by Cell Ranger. The data was analysed using the pipeline described above. Only the clusters identified as fibroblasts (based on expression of Col1a1, Col1a2) were considered for comparison with the CAF clusters.

### Human skin fibroblasts

scRNA-seq data was downloaded from ArrayExpress (E-MTAB-6831)^66^. CD45-negative cells from a digested skin sample were taken from a human female and processed in a 10X Chromium machine (10X Genomics). Droplet-based sequencing data was aligned, filtered and quantified using the Cell Ranger Single-Cell Software Suite (version 1.2.0), against the GRCh38 human reference genome provided by Cell Ranger. The data was analysed using the pipeline described above. Only the clusters identified as fibroblasts (based on expression of COL1A1, COL1A2) were considered for comparison with the CAF clusters.

### Comparison of human and mouse skin fibroblasts with CAFs

To compare the mouse and human skin fibroblasts with the CAFs, a logistic regression with L2-norm regularization and a multinomial learning approach (implemented by the scikit-learn function LogisticRegression) was trained on the CAF clusters, using the log-transformed normalized data. The model was used to predict the probabilities of each mouse and human skin cell belonging to each one of the CAF clusters (implemented by the predict_proba function).

#### Flow Cytometry

Following a 20min incubation with a fixable fluorescent viability stain (see isolation of single cells), cells were incubated with primary antibodies, against cell surface markers, for 30mins at 4°C. All primary antibodies were diluted according to table 1 in PBS 0.5% BSA. If required, fluorescently labelled streptavidin, diluted 1:300 in PBS 0.5%BSA, was added for a further 30mins. To stain for intracellular targets samples were fixed and permeabilized using the FOXP3 kit (eBioscence), according to manufacturer’s instructions. Fixation and permeabilization was only performed once staining for surface markers was completed. For investigation of CXCL12 expression, samples were incubated with Brefeldin-A (BFA, Biolegend) prior to the staining process. Samples were incubated with BFA (1:1000 dilution) both during tissue processing and for a further 4hs in 10% FBS supplemented DMEM media (Life Technologies). Once staining was completed, samples were analysed using the BD LSR-Fortessa system.

**Table 1.** Conjugated Antibodies.

| ***Target*** | ***Species*** | ***Company*** | ***Clone*** | ***Dilution*** |
| --- | --- | --- | --- | --- |
| CD45 APC-Cy7 | Rat | Biolegend | 30-F11 | 1:300 |
| CD45 FITC | Rat | Biolegend | 30-F11 | 1:300 |
| CD31 PE-Cy7 | Rat | eBioscience | 390 | 1:300 |
| PDPN APC | Syrian Hamster | Biolegend | 8.1.1 | 1:300 |
| CD3e 488 | Armenian Hamster | Biolegend | 145-2C1 | 1:300 |
| CD3e PE | Armenian Hamster | Biolegend | 145-2C1 | 1:300 |
| CD4 PE-Cy7 | Rat | eBioscience | GK1.5 | 1:300 |
| CD8 780 | Rat | eBioscience | 53-6.7 | 1:300 |
| CD8 PE | Rat | eBioscience | 53-6.7 | 1:300 |
| Lag3 Biotin | Rat | Biolegend | C9B7W | 1:300 |
| PD1 | Rat | Biolegend | RMP1-30 | 1:300 |
| Ki67 | Rat | Biolegened | 16A8 | 1:100 |
| IL-7Ra APC | Rat | Biolegend | A7R34 | 1:300 |
| B220 488 | Rat | Biolegend | RA3-6B2 | 1:300 |
| CD11b 647 | Rat | Biolegend | M1/70 | 1:300 |
| CD11c PE-Cy7 | Armenian Hamster | Biolegend | N418 | 1:300 |
| aSMA | Mouse | Thermo Fisher | 1A4 | 1:200 |
| PDGFRa Biotin | Rat | Biolegend | APA5 | 1:300 |
| PDGFRb Biotin | Rat | Biolegend | APB5 | 1:300 |
| Thy1 APC-Cy7 | Rat | Biolegend | 30-H12 | 1:300 |
| CD34 APC | Armenian Hamster | Biolegend | HM34 | 1:200 |
| CXCL12 PE | Mouse | R&D Systems | MAB350 | 1:100 |

### Immunofluorescence

Collected tissues were embedded in OCT medium (VWR) and snap frozen on dry ice. Frozen tissues were sectioned into 10μm slices and stored at −80°C. Sections were air dried and fixed in 50:50 acetone (Fluka): methanol (Fisher), at −20°C for 2mins or 4% paraformaldehyde (PFA) for 10 minutes. If fixed with PFA, samples were permeabilized with 0.1 % Triton for a further 10 minutes. After blocking for 1h at room temperature (RT) with blocking buffer (10% chicken serum and 2% Bovine Serum Albumin) in PBS, primary antibodies were applied overnight at 4°C or RT for 3hs. To visualise samples, secondary antibodies (Life Technologies), conjugated to either Alexa Fluor 488, 594 or 647, or fluorescently labelled streptavidin, were added for 2hs at RT. Samples were incubated with the nuclear stain 4′,6-diamidino-2-phenylindole (DAPI) for 10 mins at 1μg/ml, before mounting with ProLong Gold (ThermoFisher) liquid mountant. Streptavidin and secondary antibodies were diluted 1:300 in blocking buffer and primary antibodies were diluted in blocking buffer according to Table 2. Confocal imaging was performed using the Zeiss LSM 880 microscope and processed using the Zeiss Blue software.

**Table 2.** Purified antibodies.

| ***Target*** | ***Species*** | ***Company*** | ***Clone*** | ***Dilution*** |
| --- | --- | --- | --- | --- |
| PDPN | Syrian Hamster | Biolegend | 8.1.1 | 1:100 |
| αSMA | Rabbit | abcam | Polyclonal | 1:50 |
| PDGFRα | Goat | R&D Systems | Polyclonal | 1:50 |
| CD34 | Rat | eBioscience | RAM34 | 1:50 |
| F4/80 | Rat | AbDserotech | A3-1 | 1:100 |
| F4/80 488 | Rat | AbDserotech | A3-1 | 1:20 |
| CXCR4 | Rat | R & D Systems | 247506 | 1:50 |
| CSFR1 | Sheep | R & D Systems | Polyclonal | 1:50 |
| CXCL12 PE | Mouse | R & D Systems | MAB350 | 1:50 |
| CSF1 | Rabbit | ABGENT | Polyclonal | 1:50 |
| NG2 | Rabbit | abcam | Polyclonal | 1:50 |
| CD31 | Rat | Biolegend | MEC13.3 | 1:100 |

### EdU Incorporation

B16 melanomas were established in wt C57BL/6 mice as previously stated. Tumours were collected after 11 days and frozen in OCT medium for histology. Intraperitoneal injections of 500μg/ml of 5-ethynyl-2’-deoxyuridine (EdU) were performed every 24hs, 4 days prior to culling. Sections were fixed at −20°C, in a mixture of acetone and methanol (50:50). After fixation, the EdU Click-it Alexa Fluor 647 kit (Invitrogen) was used to visualise incorporated EdU, according to the manufacturer’s protocol. Following the click-it reaction, antibody staining was performed as previously stated.

### Data availability

The raw sequencing data for the melanoma model has been deposited in ArrayExpress (experiment E-MTAB-7427) and the count table can be downloaded from https://www.ebi.ac.uk/gxa/sc/experiments/E-EHCA-2/Results. The mouse skin data from healthy mice was deposited in ArrayExpress (experiment E-MTAB-7417).

## Acknowledgments

We thank all members of Teichmann and Shields Lab for helpful discussions.

## Author contributions

S.A.T, J.D.S and B.M conceived the study. S.D, A.R, B.M and J.P performed the mouse experiments, sample/library preparation and histology staining. M.E and S.D analysed the data and interpreted the results with contributions from J.H and G.K. T.H and X.C performed the sample/library preparation for the healthy mouse skin samples. S.D, M.E, J.D.S and S.A.T wrote the manuscript. S.A.T and J.D.S co-directed the study. All authors read and accepted the manuscript.

## Extended Data Figures

**Extended Data Figure 1. Gating strategy and quality control of SS2 data. a**, Gating strategy for index sorted cell populations. **b**, Quality control of the scRNA-seq dataset. Histograms show distribution of the cells from all cells that passed the computational quality control ordered by number of detected genes and mitochondrial gene expression content.

**Extended Data Figure 2. Marker genes**. **a**, Heatmap showing relative expression (z-score) of the top 5 markers for each cluster presented in Fig. 1B.

**Extended Data Figure 3. Overview of the innate subpopulations. a**, tSNE plots visualisation of expression of MHC class I and II genes in the innate immune subpopulations. **b**, Violin plots showing expression of selected cytokines for each innate immune subpopulation, known to induce T-cell responses. **c**, Heatmap of gene ontology pathways for differentially expressed genes in innate immune subpopulations; Antigen presentation and cytokine-chemokine receptor interactions. Columns represent individual cells, rows display z scores. d, Bar plot (-log FDR) depicting the top 20 gene ontology pathways upregulated in cDC1 cells, located in the tumour compared to the lymph node (left). Heat map depicting genes in the Class 1 MHC mediated antigen presentation pathway (right). Columns represent individual cells, rows display z scores.

**Extended Data Figure 4. Macrophage markers. a**, tSNE plots showing expression of M1 and M2 macrophage markers in the innate immune subpopulations.

**Extended Data Figure 5. a**, Percentage of CD8^+^ T cells (displayed as percentage of total T cells) and percentage of Ki67^+^, Lag3^+^ and PD1^+^, T cells (displayed as a percentage of total CD8^+^ T cells), in tumours and lymph nodes isolated from skin, day 5 and day 11 tumour bearing or control mice. Data presented as mean ± SEM. * P<0.05 as calculated using either a one way anova with Tukey post-hok test, n=4 independent mice per time point.

**Extended Data Figure 6. Distinct CAF subpopulations identified in the melanoma mouse model. a**, tSNE plots showing expression of typical CAF markers. **b**, Bar plot (-log FDR) depicting the top 50 gene ontology pathways upregulated in each CAF population. C, IF imaging of EdU incorporation in a subset of CAFs in day 11 tumours. Scale bars 50μm **(c, d**, Representative confocal images of CSF1 and CXCL12 mRNA transcripts in CD34+ PDGFRα+ fibroblasts of day 11 tumours following RNAscope *in situ* hybridization technology. Scale bars 20um.

**Extended Data Figure 7. a**, Heatmap showing expression of canonical fibroblasts and pericytes markers in the CAFs. **b**, tSNE plots of all sequenced cells, showing expression of typical pericyte markers is also detected in PDPN^+^ lymph node fibroblasts. **c** and **d**, IF imaging showing αSMA^+^ and NG2^+^ cells both distinct from (top panel) and associated with

(middle panel) CD31^+^ blood vessels. Bottom panels show an abundance of CD31^+^ blood vessels that are not surrounded by pericytes.

**Extended Data Figure 8. CAF validation and human clustering. a**, Gating strategy for flow cytometry characterization of CAFs. **b**, tSNE plots showing expression of CAF marker genes in the human melanoma dataset^24^.

## Supplementary Tables

**Supplementary table 1. Differentially expressed genes amongst the innate immune system clusters.** This table provides the differentially expressed genes between the innate immune subpopulations (*q*-value<0.1).

**Supplementary table 2. Pseudotime analysis of the CD8 T cells.** Genes identified as varying significantly along the CD8 T cells trajectory (*q*-value<0.1).

**Supplementary table 3**. **Differentially expressed genes amongst the three CAF clusters.** This table provides the differentially expressed genes between the three CAF subpopulations (*q*-value<0.1).

**Supplementary table 4. List of interactions in the tumour infiltrating cells.** This table provides the list of interaction pairs between clusters in Fig. 1B resulting from our cell-cell communication pipeline. Sheet 1 - *p*-values; Sheet 2 - mean expression of the average receptor expression level of a cluster and the average ligand expression level of the interacting cluster; Sheet 3 - mean expression of the significant interactions ranked by specificity (*p*-value < 0.05).

